# Development of a Robust Consensus Modeling Approach for Identifying Cellular and Media Metabolites Predictive of Mesenchymal Stromal Cell Potency

**DOI:** 10.1101/2023.02.03.526990

**Authors:** Alexandria Van Grouw, Maxwell B. Colonna, Ty S. Maughon, Xunan Shen, Andrew M. Larey, Samuel G. Moore, Carolyn Yeago, Facundo M. Fernández, Arthur S. Edison, Steven L. Stice, Annie C. Bowles-Welch, Ross A. Marklein

**Affiliations:** School of Chemistry and Biochemistry and Parker H. Petit Institute for Bioengineering and Bioscience, Georgia Institute of Technology, Atlanta, Georgia, USA; Department of Biochemistry & Molecular Biology, Complex Carbohydrate Research Center and Institute of Bioinformatics, University of Georgia, Athens, Georgia, USA; School of Chemical, Materials, and Biomedical Engineering, Regenerative Bioscience Center, University of Georgia, Athens, Georgia, USA; Marcus Center for Therapeutic Cell Characterization and Manufacturing, Parker H. Petit Institute for Bioengineering and Bioscience, Georgia Institute of Technology, Atlanta, Georgia, USA; Regenerative Bioscience Center, Department of Animal and Dairy Sciences, University of Georgia, Athens, Georgia, USA; Systems Mass Spectrometry Core, Parker H. Petit Institute for Bioengineering and Bioscience, Georgia Institute of Technology, Atlanta, Georgia, USA

**Author notes:** These authors contributed equally (Alexandria Van Grouw, Maxwell B. Colonna, Ty S. Maughon). **Contact Information:** Ross Marklein, 220 Riverbend Rd, Athens, GA 30602, Annie C. Bowles-Welch, 345 Ferst Dr. NW, Atlanta, GA 30318. **Author contributions:** AVG: Collection and assembly of data, data analysis and interpretation, manuscript writing MBC: Collection and assembly of data, data analysis and interpretation, manuscript writing TSM: Conception and design, collection and assembly of data, data analysis and interpretation, manuscript writing XS: Collection and assembly of data AML: Collection and assembly of data SGM: Collection and assembly of data CY: Conception and design, financial support FMF: Conception and design, financial support, manuscript writing and final approval of manuscript ASE: Conception and design, financial support, manuscript writing and final approval of manuscript SLS: Conception and design, financial support, manuscript writing and final approval of manuscript ABW: Conception and design, collection and assembly of data, data analysis and interpretation, manuscript writing, final approval of manuscript RAM: Conception and design, data analysis and interpretation, financial support, manuscript writing and final approval of manuscript.

**Keywords:** MSCs, machine learning, immunomodulation, critical quality attributes, metabolism

## Abstract

Mesenchymal stromal cells (MSCs) have shown promise in regenerative medicine applications due in part to their ability to modulate immune cells. However, MSCs demonstrate significant functional heterogeneity in terms of their immunomodulatory function because of differences in MSC donor/tissue source, as well as non-standardized manufacturing approaches. As MSC metabolism plays a critical role in their ability to expand to therapeutic numbers *ex vivo*, we comprehensively profiled intracellular and extracellular metabolites throughout the expansion process to identify predictors of immunomodulatory function (T cell modulation and indoleamine-2,3-dehydrogenase (IDO) activity). Here, we profiled media metabolites in a non-destructive manner through daily sampling and nuclear magnetic resonance (NMR), as well as MSC intracellular metabolites at the end of expansion using mass spectrometry (MS). Using a robust consensus machine learning approach, we were able to identify panels of metabolites predictive of MSC immunomodulatory function for 10 independent MSC lines. This approach consisted of identifying metabolites in 2 or more machine learning models and then building consensus models based on these consensus metabolite panels. Consensus intracellular metabolites with high predictive value included multiple lipid classes (such as phosphatidylcholines, phosphatidylethanolamines, and sphingomyelins) while consensus media metabolites included proline, phenylalanine, and pyruvate. Pathway enrichment identified metabolic pathways significantly associated with MSC function such as sphingolipid signaling and metabolism, arginine and proline metabolism, and autophagy. Overall, this work establishes a generalizable framework for identifying consensus predictive metabolites that predict MSC function, as well as guiding future MSC manufacturing efforts through identification of high potency MSC lines and metabolic engineering.

## INTRODUCTION

Preclinical and clinical studies have demonstrated evidence of promising therapeutic effects from the use of mesenchymal stromal cells (MSCs) for a broad range of applications including autoimmune, neurodegenerative, and inflammatory diseases.^1–3^ MSCs are derived from various tissues of the body, most commonly bone marrow, adipose, and umbilical cord,^4–7^ and can be administered directly as cell therapies or used to create cell-derived products (e.g., secretome and extracellular vesicles).^8–10^ The secretory repertoire of MSCs is rich in cytokines, chemokines and growth factors that, combined with the fact that MSCs lack or possess low expression of self-antigens (thus permitting allogeneic use), renders MSCs a potential cell therapy for a large patient population with life-altering conditions.^9,10^ However, efficacy outcomes from clinical trials are inconsistent and pose a major roadblock to the approval of MSCs as cell therapies despite a well-established safety record.^11^ One major challenge that must be addressed to facilitate clinical translation is MSC functional heterogeneity, which can be attributed to different donors, tissue sources and culture conditions (i.e., ‘manufacturing’) introduced during MSC *ex vivo* expansion.^1,12,13^ MSCs require *ex vivo* expansion to yield a sufficient cell supply to meet the needs for clinical dosages, which are typically on the order of hundreds of millions of cells.^14^ Relatedly, a lack of standard cell culture practices and processes is another challenge that contributes to poor reproducibility and inconsistent clinical outcomes.^15^ Better understanding of MSC functional heterogeneity will lead to critical quality attributes (CQAs), which are part of a comprehensive analytical assay suite to be used for MSC release criteria, ultimately improving their clinical translation.

CQAs predictive of MSC immunomodulatory potency serve as a quantitative and reproducible measure for assessing MSC quality, thus enabling rational approaches to both identify high quality MSC lines (donors), as well as optimize culture conditions that facilitate scaling to larger manufacturing formats such as bioreactors. Several approaches to identify CQAs correlative to MSC functional potency have been reported.^16,17^ Based on previous work ^10,18^, MSC potency assays often assess MSC immunomodulatory functions such as indoleamine 2.3-dioxygenase (IDO) activity, as well as MSC modulation of activated immune cells (e.g. T cells and macrophages). These functions are relevant to a broad range of diseases and have been associated with MSC secretion of anti-inflammatory, mitogenic, and tissue reparative factors.^19,20,19,20,21,22^ priming MSCs with stimulatory molecules (e.g., interferon-γ (IFN-γ) or tumor necrosis factor-α (TNF-α)) promotes the increased secretion of paracrine mediators such as IDO and anti-inflammatory molecules (e.g., interleukin-10) that are relevant in the context of MSC immunomodulation (e.g., suppress T cell proliferation, induce regulatory phenotypes of T cells and macrophages).^21,22^ Not only can these MSC-mediated effects related to immunomodulatory potency establish a set of CQAs, they shed light on metabolic activities that are vital to the basic understanding of MSC functions that can be leveraged for therapeutic use.^23^ However, the reproducibility and robustness of these potency assay formats reduce their appeal for assessing CQAs in a manufacturing setting.

Robust multi-omic approaches coupled with computational modeling to identify correlations to functional potency have been leveraged to identify predictive markers in cell therapies.^24–26^ Metabolomics is an emerging field due to the abundance of metabolites and their reflection of the cellular phenotype which allows for non-targeted approaches for important biomarker discovery.^27,28^ Well established techniques such as nuclear magnetic resonance (NMR) and mass spectrometry (MS) are often used to measure both cellular and extracellular metabolites.^29 30^ In terms of MSC metabolism, it has been shown that MSCs preferentially utilize glycolysis over oxidative phosphorylation (OXPHOS) *in vivo*, but shift towards OXPHOS from extended culture and expansion.^31^ Greater OXPHOS metabolism has also been shown to lead to a decrease in T cell suppression by MSCs.^23^ Previously, our group assessed end of expansion intracellular metabolites as candidate predictive markers, and putative CQAs, using three independent MSC lines at three passages. Partial least squares regression (PLSR) was used to determine predictive markers based on their variable importance projection (VIP) score.^26^ Several amino acids, small molecule metabolites (e.g. myo-inositol), and phosphatidylcholines (PCs) were shown to be correlative to MSC functional potency (measured in terms of both T cell modulation and IDO activity).

In this study, we expanded on our previous work by applying a novel CQA discovery framework whereby metabolomic data generated both in-process (during the first three days of MSC expansions) and at end of production (intracellular metabolites from harvested MSCs) were input into a suite of models to predict MSC immunomodulatory potency. For this study, we generated ten MSC lines consisting of the following: the same three cell lines as our previous study, six additional MSC lines, and a repeat expansion of one of the MSC lines.^26^ For each MSC line, unsupervised analysis of media metabolites and intracellular metabolome of MSCs were correlated to a functional composite score (based off T cell suppression and IDO activity) using several machine learning (ML) models. CQAs were resolved across a total of 7 models (for both extracellular and intracellular inputs) to identify consensus metabolites that were subsequently related using metabolic pathway analysis. Top consensus media metabolites predictive of MSC potency include several amino acids (proline, arginine, aspartate), pyruvate and fructose. Top consensus intracellular metabolites predictive of MSC potency include several lipid classes such as PCs, phosphatidylethanolamines (PEs), and sphingomyelins (SMs). These were then mapped to several metabolic pathways to infer their potential biological roles as they relate to MSC immunomodulation. This study establishes a generalizable framework for identifying predictive markers from early stage culture media and intracellular metabolomic analyses that can ultimately be used to select for MSC lines with desired properties and guide future MSC manufacturing strategies.

## MATERIALS AND METHODS

### MSC Expansion and MSC/MSC-Conditioned Medium Sample Preparation

Bone marrow-derived MSCs (BMMSCs) were purchased from RoosterBio, Inc. (Frederick, MD), and iMSCs were purchased from Fujifilm Cellular Dynamics Inc (Madison, WI). Prior to this study’s expansion, MSCs were previously expanded to an initial population doubling level (PDL_0_) reported in **Supporting Information Table 1**. Additional information for each MSC cell line including final PDL, donor demographic information, and final cell yield is also reported in this table. Cryopreserved vials from each donor were thawed, and 10^6^ MSCs were seeded into an initial T-150 tissue culture flask in complete media containing Gibco™ Minimum Essential Media α with nucleosides (Thermo Fisher Scientific, Waltham, MA), 10% fetal bovine serum (FBS; HyClone Laboratories, Logan, UT), and 1% penicillin-streptomycin solution (10,000 U/mL; Sigma-Aldrich, St. Louis, MO) for a culture rescue period of 48 hr. The same lot of FBS was used throughout the study. MSCs were then washed with endotoxin-free Dulbecco’s phosphate buffered saline (PBS) without calcium and magnesium (Millipore Sigma), harvested using 1X TrypLE™ Express Enzyme (Thermo Fisher Scientific), neutralized with complete media, and centrifuged 300g to create a cell pellet. MSCs were then resuspended in complete media and counted. Next, MSCs from each donor were seeded at 500 cells/cm^2^ into 10 T-75 flasks containing 10 mL complete media. Control flasks containing 10 mL complete media only were also prepared. All flasks were then transferred to a humidified incubator set to 37° C and 5% CO_2_.

MSC conditioned medium (CM) sample collection of 300 μl was performed for each flask at approximately the same time each day (±1 hr) and total complete media exchange was performed every 3 days until MSCs achieved 70-80% confluence. All media samples were placed directly into −80° C storage until further analysis by NMR. MSCs were then harvested using the same procedure described above. Cell pellets were split for cryopreservation (and functional analysis, see below) or preparation for intracellular lipidomic/metabolic analysis. Cell pellets designated for cryopreservation were prepared into cryovials containing 10^6^ MSCs in 1 mL CryoStor^®^ CS 10 (Sigma-Aldrich) and stored at −80° C for 24 hr using controlled rate freezing containers. Vials were then transferred to the vapor phase in a liquid nitrogen cryogenic freezer until further analysis. For intracellular lipidomic/metabolomic analysis, cell pellets were washed twice by resuspending in PBS and centrifuged at 10,000 rpm. All supernatant was removed and cell pellets were then stored at −80° C.

### MSC Functional Analysis - T cell Suppression Assay

MSCs from each cell-line were thawed and allowed to recover for 48 hours with a media change at 24 hours. MSCs were harvested using TrypLE then seeded at a density of 10,000 cells/well in a 96 well plate and cultured for 24 hours. Previously frozen peripheral blood mononuclear cells (PBMCs, AllCells, Alameda CA) were thawed in RPMI media (RPMI, 20% FBS, 2mM L-glutamine, 50 U/mL penicillin, 50 μg/mL streptomycin) and cultured overnight at 37°C and 5% CO2. Prior to co-culture, PBMCs were labeled with CFSE (**Supporting Information Table 2**, Biolegend, San Diego CA) according to the manufacturer’s protocol, and 100,000 PBMCs were added to each well at a final MSC:PBMC ratio of 1:10 as well as control wells containing only PBMCs. Following PBMC addition, stimulating anti-CD3/CD28 Dynabeads (Thermo Fisher Scientific, Waltham MA) were added at 100,000 beads per well to the appropriate wells (positive controls and all MSC groups). MSCs and PBMCs were co-cultured for 72 hours at 37°C, 5% CO_2_. Following co-culture, PBMCs were collected and stained using Brilliant Violet 711 anti-CD4 and APC anti-CD8 (**Supporting Information Table 2**) (Biolegend, San Diego CA). PBMCs were first washed and stained with Zombie Yellow (**Supporting Information Table 2**) (Biolegend, San Diego CA) viability dye and blocked using 2% FBS. PBMCs were then washed again and stained for CD4 and CD8 in the dark at room temperature. Following staining, the antibodies were blocked using 2% FBS and washed. PBMCs were then fixed with 4% PFA for 30 minutes at 4°C, washed and re-suspended in PBS containing 2% FBS, and finally stored overnight at 4°C in the dark until flow analysis.

All flow cytometry experiments were performed using a Novocyte Quanteon flow cytometer (Agilent, Santa Clara CA) with 20,000 events collected per sample. All flow cytometry data were analyzed using FlowJo (Treestar, Inc., Ashland OR). Briefly, cell debris, doublets, and Dynabeads were gated out using scatter principles. Then, single stained controls were used for compensation, and fluorescence minus one controls were used to determine positive populations (**Supporting Information Fig. S1**).

### MSC Functional Analysis - Indoleamine 2,3 Dioxygenase (IDO) Activity Assay

MSCs from each cell-line were thawed and cultured for 48 hours with a media change at 24 hours. MSCs were harvested using TryplE then seeded at a density of 40,000 cells/cm^2^ in a 96 well plate. After 24 hours, the medium was replaced in each well with complete medium containing 10 ng/mL IFN-γ (Life Technologies). After an additional 24 hours, conditioned medium was collected and frozen at −20°C, and cells were fixed with 4% PFA. Each medium sample was thawed and 100 μL transferred into a 96 well plate. Trichloroacetic acid was used to precipitate excess protein. 75 μL of the supernatant was collected and transferred to a separate 96 well plate. Ehrlich’s Reagent was then added to each well to detect L-kynurenine levels using a SpectraMax iD5 (Molecular Devices) plate reader. Levels of L-kynurenine were determined using a standard curve. To normalize L-kynurenine values to cell numbers for each replicate well (n=5 wells) within an experimental group, we performed automated image analysis to quantify cell nuclei in the wells from which conditioned medium was collected. Following fixation, MSCs were washed with PBS twice, and stained with Hoechst (10μg/mL) in 0.1% Tween20 PBS solution for 1hr. Cells were then imaged on a Cytation 5 high content imaging system (Biotek, Winooski VT) and cell counts determined using CellProfiler to normalize the amount of L-kynurenine per cell.

### Generation of a Composite Score of MSC Potency

Using the results from functional assays, a composite functional score was generated to use as an overall potency metric. The averages of the percentage of CFSE dilution of CD4 and CD8 T cells from two PBMC donors and IDO activity were subject to principal component analysis (PCA) using JMP software (V9.2). The first principal component (PC1) value of each cell line was taken as the respective composite functional score and represented 74.5% of the total variance of the data.

### NMR Media Analysis of MSC Culture

Frozen media samples were transported on dry ice for NMR analysis. Samples were prepared in batches for each rack of 96 1.7 mm SampleJet NMR tubes (Bruker BioSpin). Briefly, samples were pulled and sorted on dry ice, then thawed at 4°C. A solution of 3.33 mM DSS-D6 in deuterium oxide (Cambridge Isotope Laboratories) were mixed into media samples to a final concentration of 10% deuterium oxide. Samples were then centrifuged @ 2990 g for 20 min at 4°C to pellet any debris that may have been collected with the media. 60 μl were transferred from each sample tube to NMR tubes using Bruker SamplePro liquid handling system (Bruker BioSpin). Ten μl of remaining volume from each sample in the rack was combined to create an internal pool used within each rack. These internal pooled samples were placed throughout the rack as quality control during data acquisition.

NMR spectra were collected on a Bruker Avance NEO spectrometer at 800 MHz using a 1.7 mm TCI cryogenic probe and TopSpin software (Bruker BioSpin). One-dimensional spectra were collected on all samples using the noesypr1d pulse sequence under automation using IconNMR software. After one-dimensional data acquisition of all samples, 10 samples of each media type were selected at random, material collected from the NMR tubes, and combined to form a media specific pool used for two-dimensional (2D) spectral acquisition. 2D HSQC and TOCSY spectra were collected on pooled samples for metabolite annotation. One-dimensional spectra were automatically phase and baseline corrected using a batch script executed in MestreNova software (MestreLab Research). 2D spectra were processed in NMRpipe.^32^

One dimensional spectra were referenced, water/end regions removed, aligned and normalized using an in-house MATLAB (The MathWorks, Inc.) toolbox established by Robinette, et. al and implemented within NMRbox.^33,34^ Spectral features that were sufficiently well aligned across spectra were semi-automatically integrated using an interactive MATLAB function to obtain feature intensity values. These feature intensity values were batch corrected across the two batches of data collection using the ComBat algorithm^35^ implemented in Metaboanalyst^36^. The feature values for the first timepoint of each replicate was subtracted from the values of the second and third timepoints, to produce features representing the differences between them. To reduce the total number of feature differences used for machine learning analyses, a variance filter was applied to select a subset of spectral features with highest variance as previously described.^25^ For each timepoint differences being assessed, the top percentage of variable features was determined by the subset which provided the best partial least squares regression R^2^ when used as input. This resulted in 69 features used for the features for Day 3 minus Day 1 values, and 21 features for Day 2 minus Day 1 values.

### Intracellular Metabolite Analysis of MSCs Using LC-MS

Approximately one million MSCs were analyzed for each sample. Frozen cell pellets were thawed and washed prior to undergoing a modified Bligh-Dyer extraction to yield two phases. Extraction solvent (2:2:1 chloroform:methanol:water) and glass beads (400-600 μm) were added to cell pellets for extraction and homogenization in a TissueLyser II to 30 Hz for 6 minutes. Samples were then sonicated and centrifuged. Following extraction, 300 μL aliquots from each layer were transferred to new microcentrifuge tubes and solvent was dried using vacuum centrifugation. Dried organic phase samples were re-constituted in isopropyl alcohol, while dried aqueous phase samples were re-constituted in 80% methanol. Re-constitution was followed by sonication, centrifugation, and transfer to liquid chromatography (LC) vials for ultrahigh performance liquid chromatography mass spectrometry (UHPLC-MS) analysis (**Supporting Information Table 3**). Media samples without cells were also analyzed as blanks to remove any features corresponding to remaining media components on the cells. Ten μL of media was subject to the same Bligh-Dyer extraction as above and extracts were run according to the instrumental methods listed above. A quality control (QC) sample for hydrophilic interaction chromatography (HILIC) and reverse phase datasets was created by pooling 20 μL from each experimental sample. The pooled QC injections were used for drift correction of peak areas. Sample queue was randomized with a mix of samples, QCs, and blanks.

Metabolomic analysis was performed on all aqueous extracts using UHPLC-MS on an Orbitrap ID-X Tribrid mass spectrometer (ThermoFisher Scientific). HILIC separation was employed with a Waters ACQUITY UPLC BEH amide column (2.1 ×150 mm, 1.7 μm particle size) on a Vanquish (ThermoFisher Scientific) chromatograph. Lipidomic analysis was performed on all organic extracts using UHPLC-MS on a Q Exactive HF Hybrid Quadrupole-Orbitrap mass spectrometer system (ThermoFisher Scientific). Reverse Phase separation was employed with an Accucore™ C30 column (2.1×150 mm, 2.6 μm particle size) on a Vanquish (ThermoFisher Scientific) chromatograph. Data dependent acquisition (DDA) was employed in both instrument methods to yield fragmentation information for detected metabolites and lipids. DDA methods collected full scan data at resolution of 120,000 in the orbitrap. This was followed by collection of fragmentation spectra (MS^2^) of selected precursors collected in the ion trap with an isolation window of 0.4 *m/z* with a cycle time of 1.25 s. Dynamic exclusion was set to exclude MS^2^ collection of precursors within a 6 s window with a 10 ppm mass tolerance for the precursor ion. Stepped normalized collision energies of 15%, 30%, and 45% were employed with HCD and CID 35% to collect MS^2^ spectra.

Data was preprocessed in Compound Discoverer 3.2 (Thermo Fisher Scientific). This included drift correction and blank removal. All features with at least 5× the signal in the blanks and were present in at least 50% of all pooled QC injections were kept for analysis. This resulted in a dataset with a total of 8,388 features. Further data filtering was used to remove features with relative standard deviations lower than 25% between all samples, features with poor chromatographic peak shape, and features with no MS^2^ spectral data. Because the experimental samples were prepared at two distinct times, a batch correction (**Supporting Information Fig. S2**) was employed to harmonize the data. Feature peak areas were corrected across the two batches using the ComBat algorithm^3,4^ implemented in Metaboanalyst^3,5^. Annotations were performed based on exact mass matches, MS^2^ spectra fragmentation patterns, and MS^2^ spectral library database matches. The final datasets used for machine learning included 436 annotated lipids and 43 annotated small polar analytes. The peak areas in these datasets were median normalized and autoscaled prior to machine learning workflow.

### Computational Analysis and Statistical Methods

Identification of features predictive of the functional composite score across MSC donors was based on methods described previously^25^ using multiple machine learning methods. In brief, the ML regression methods utilized were random forest (RF), gradient boosted regression (GBR), decision tree regression (DTR), least absolute shrinkage and selection operator (LASSO), partial-least squares regression (PLSR), support vector regression (SVR), and symbolic regression (SR). These models were used to extract predictive variables (or variable combinations). SR was performed using Evolved Analytics’ Data Modeler software (Evolved Analytics LLC). All other models were generated with the LinearSVR, PLSRegression, RandomForestRegressor, DecisionTreeRegressor, Lasso, and GradientBoostingRegressor functions as part of the sklearn software package implemented in Python.^37^ Parameter tuning was done for all sklearn models in a grid search manner using the GridSearchCV function with 5-fold cross validation (CV) and using R^2^ as the scoring criteria. For each regression model, feature selection was performed using the same regression type (i.e., LASSO). Final prediction performance was measured by calculating leave-one-out R^2^ (LOO-R^2^) values on final models with CV-optimized parameters. Model specific parameters and parameter ranges that were used are available in code.

Consensus analysis of the relevant variables extracted from each ML model was done to identify consistent predictive features of function using both in-process media features (measured by NMR) and end-product cellular lipids and metabolites (measured by LC-MS). For RF, GBR, DTR, LASSO, PLSR, SVR models, features that ranked in the top 20% of feature importance were selected, while for SR variables present in ≥10% of the top-performing SR models from Data Modeler (R^2^ ≥ 90%, complexity ≤300) were chosen to investigate consensus. Those variables that appeared as important in 2 or more ML methods were deemed consensus features and included for further annotation (NMR) and pathway analysis (NMR and MS).

Because the iMSC sample was the only non-bone marrow derived line, this could be an outlier. To determine the importance of the iMSC sample on the models, the final metabolite panels in the consensus models were used to create models without the iMSC sample. All modeling parameters were kept the same with the exception of changing the 5-fold cross validation to 3-fold to accommodate 9 samples. The success of these models was evaluated using LOO-R^2^. All statistics were performed in Python or Prism (GraphPad Software, San Diego CA).

### Metabolite Pathway Analysis

Enrichment analysis of consensus metabolites from both NMR and MS was performed using Metaboanalyst. Specifically, the list of consensus metabolites was submitted to perform over representation analysis by mapping to all pathways in the small molecule pathway database (SMPDB) From the list of consensus NMR metabolites, 6 out of 7 consensus metabolites were able to be mapped by Metaboanalyst, while all 16 MS consensus metabolites were mapped. Lipid pathway enrichment was performed with LIPEA, using the default background for *Homo sapiens.*16 out of 33 submitted consensus lipid species were mapped to KEGG lipids used by the LIPEA program. Final lists of mapped metabolites/lipids used for pathway enrichment are included in **Supporting Information Table 4**.^38^

## RESULTS

### Cell-line Dependent Differences in MSC Potency

Following expansion and cryopreservation of all MSC lines, MSCs were thawed to determine their functional capacity. Proliferation of CD4^+^ and CD8^+^ T cells (based on %CFSE dilution) from two PBMC donors and IDO activity (L-kynurenine production) were used to evaluate each MSC functional capacity between MSC lines and create a functional composite score (**Fig. 1A**). Similar trends were observed for CD4^+^ (**Fig. 1B, D**) and CD8+ (**Fig. 1C, E**) proliferation for both PBMC donors with MSCs suppressing T cells in a broad range from 46.8-67.4% (CD4^+^) and 54.2-77.0% (CD8^+^) for PBMC Donor 1 and 10.7-46.2% (CD4^+^) and 14.8-38.1% (CD8^+^) for PBMC Donor 2. This observation was evident by regressing CD4^+^ vs CD8^+^ T cell proliferation results for both PBMC Donor 1 and PBMC Donor 2 (R^2^=0.86 and R^2^=0.90, respectively) (**Supporting Information Fig.S3**). The iMSC line had the highest suppression of CD4^+^ and CD8^+^ T cell proliferation in both PBMC donors. RB182 was not significantly different from the iMSCs for CD4^+^ and CD8^+^ T cell proliferation in PBMC donor 1 but was for PBMC donor 2. RB71 consistently had the least amount of CD4^+^ and CD8^+^ T cell suppression from both donors. All MSC lines were significantly different from the positive control (dotted line). The MSC lines with the highest T cell suppression function (iMSC, RB177, and RB182) were also high in IDO activity (**Figure 1F**). While MSC line RB71 had the lowest observed T cell suppression, its IDO activity was not the lowest of the 10 MSC lines as its activity was approximately in the middle range (46.3 pg/cell/day) of all observed IDO activity values (20.5-80.6 pg/cell/day). During cell expansion, a repeat line, derived from RB174, was used to compare different expansion dates (i.e. batches). Functional comparison between these two expansions (termed RB174_1 and RB174_2 for batch 1 and 2, respectively) showed no significant difference within any assay. A functional composite score was generated with PCA using all the functional assay results. As PC1 comprised 74.5% of the variance in the dataset, we used PC1 values for the composite score (**Figure 1G**). This functional composite score displays a wide range of immunomodulatory function between all MSC lines with lower PC1 scores indicating MSC lines with the highest potency i.e. high T cell suppression and IDO activity.

**Figure 1.**
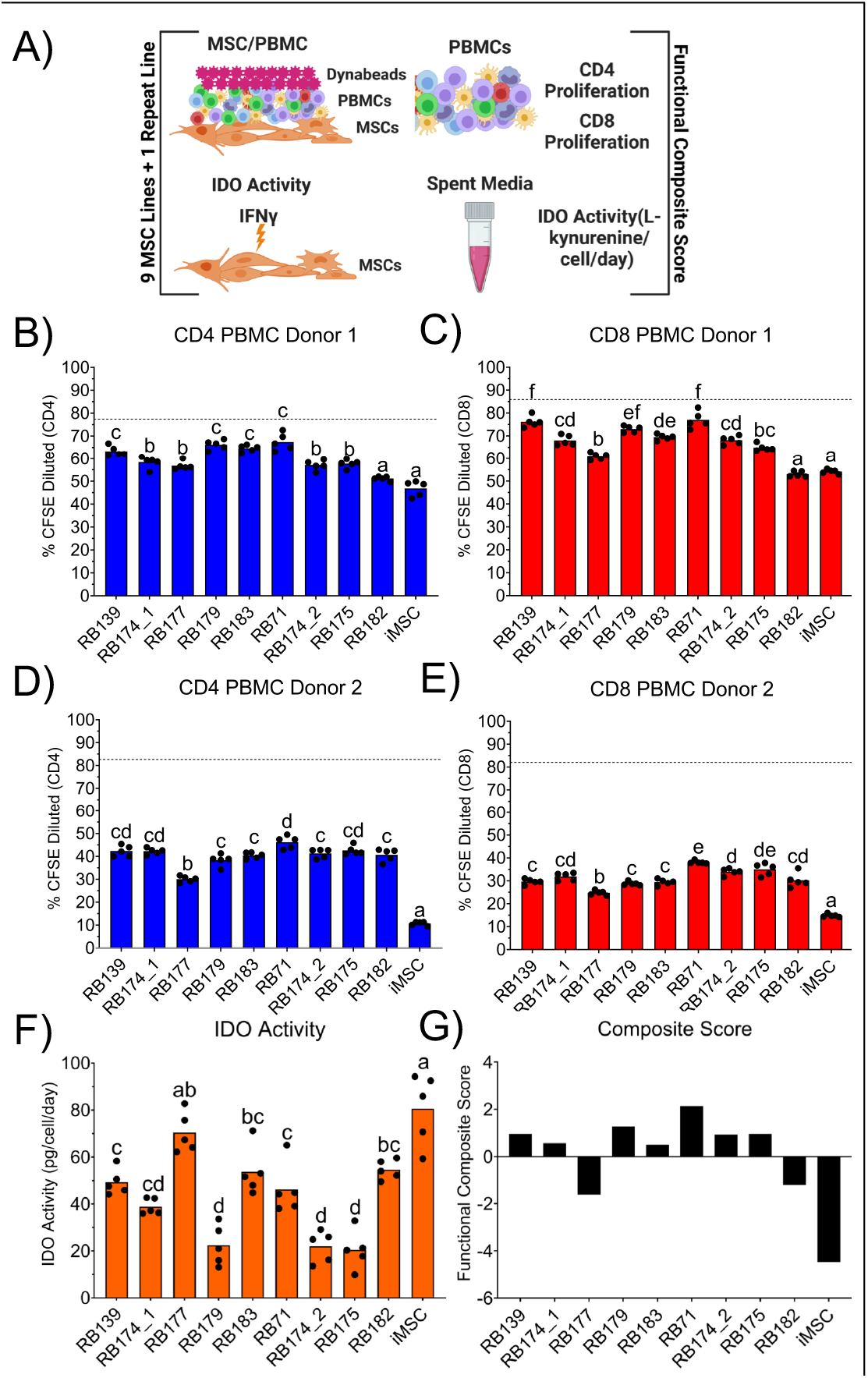
Functional analysis of IMSCs at the end of expansion. (A) Schematic of functional assays used to generate functional composite score. CD4 (B) and CD8 (C) T cell proliferation of PBMC donor 1 and CD4 (D) and CD8 (E) T cell proliferation of PBMC donor 2 based on %CFSE dilution. (F) IDO activity measured by levels of L-kynurenine in terms of pg/cell/day. (G) Functional composite score based on results of all assays (B-F). All statistics were calculated using a one-way ANOVA with Tukey’s post hoc test. Differences in letters indicate a significant difference (P<0.05) between MSC lines.

### Cell-line Differences in Intracellular MSC Metabolome

UHPLC-MS analysis by reverse phase and HILIC chromatography high-resolution mass spectrometry (**Supporting Information Table 3**) of MSC cell pellets yielded a rich metabolomic dataset with a total of 479 annotated features. Annotations were assigned using exact mass and MS^2^ spectral library matches. This feature list and the corresponding abundances were used to conduct unsupervised clustering to observe metabolic differences in the 10 MSC lines examined in this study. Sample clustering with the metabolites measured in this dataset showed that the iMSC sample had significant metabolic differences from all the bonemarrow derived cell lines using both Ward clustering (**Fig. 2A**) and PCA (**Fig. 2B**). Little clustering was observed according to potency except in the case of the middle performing cell lines, which have more metabolic similarity. However, the MSC lines with highest potency (iMSC) and lowest potency (RB71) were maximally separated based on their metabolic signatures using both Ward clustering and PCA.

**Figure 2.**
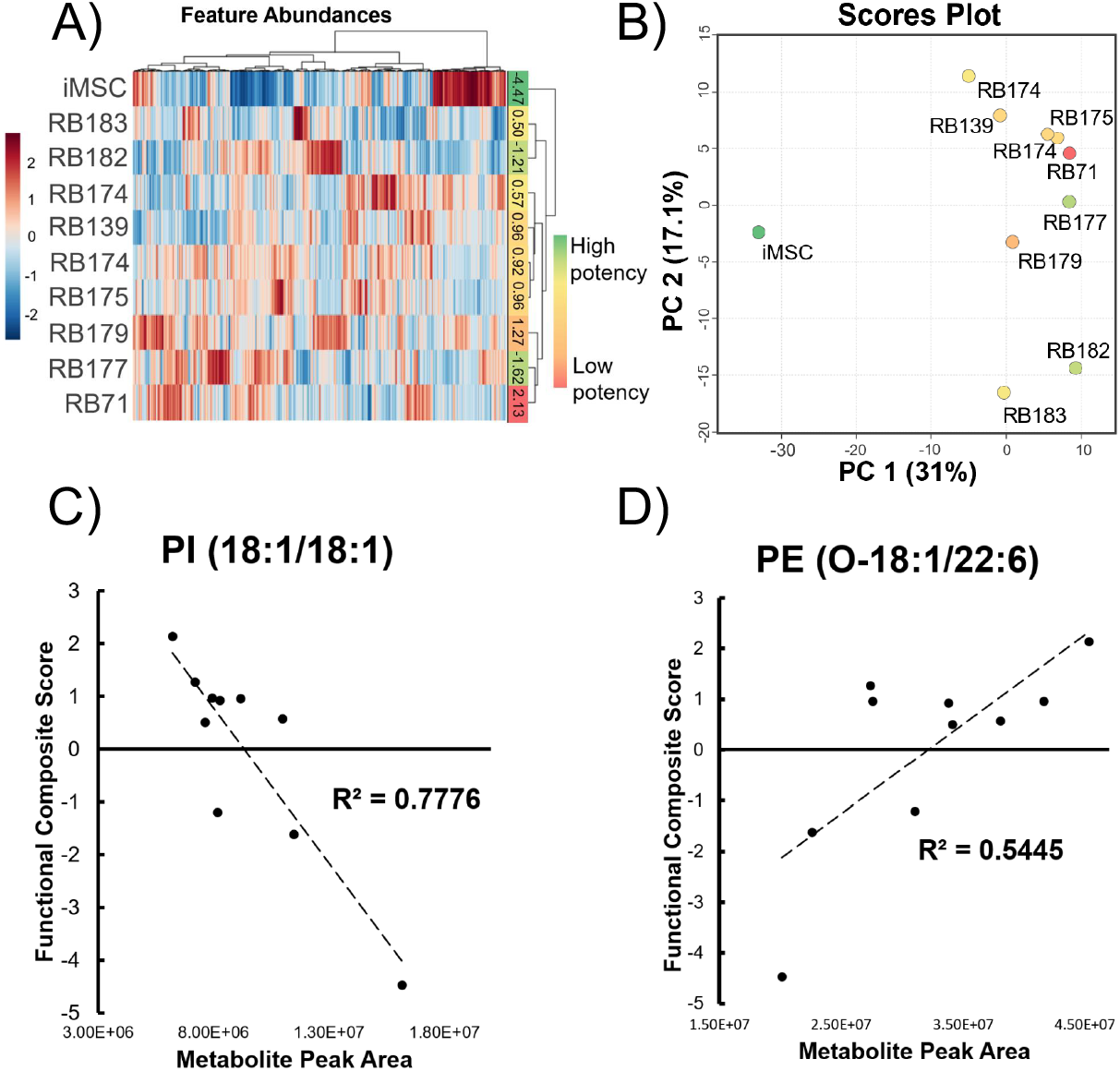
Mass spectrometry metabolomics analysis of MSC lines. Heatmap (A) and PCA scores plot (B) of ten MSC pellet samples with all 479 annotated features in the UHPLC-MS dataset that fed into the ML regression workflow. Samples are color coded according to the composite functional score determined from the functional assay results (Figure 1G). Red indicates a higher score (i.e. lower function) in immunomodulatory assays. Green indicates a lower score (i.e. higher function). Heatmap uses Euclidean distance measure and Ward clustering. Simple linear regressions of the abundances of lipids PI (18:1/18:1) (C) and PE (O-18:1/22:6) (D) against the functional composite score for 10 MSC samples.

We initially sought to find single metabolites that could predict MSC potency by examining linear correlations of metabolite abundances with the composite score. Simple linear regressions were made for each analyte in the dataset and ordered according to R^2^ (**Supporting Information Table 4C**). There were 18 annotated metabolites with R^2^ above 0.5 in the dataset. Two of these, PI (18:1/18:1) and PE (O-18:1/22:6), are shown in Fig. 2C and 2D, respectively. As few of the individual metabolites were predictive of MSC potency after multiple testing correction it was determined that a panel of metabolites identified through machine learning (ML) based regressive methods would yield higher predictive power, as well as elucidate possible pathways involved in the regulation of these metabolites.

### Single In-process Media NMR Features Do Not Correlate with Potency Measures

Media samples collected at daily intervals during cell expansions were analyzed by 1D ^1^H-NMR. In addition, 2D NMR spectra were collected on pooled material to aid in annotation of spectral features in the 1D spectrum. After spectral processing and alignments, a total of 138 spectral features were semi-automatically selected and quantified across all samples. These spectral features correspond to a smaller number of metabolites, each metabolite having one too many spectral features based upon its chemical structure. These features were initially left unannotated in order to not exclude features corresponding to unknown metabolites that may have useful value in downstream analyses. To avoid the assumptions of linear or monotonic relationships of feature intensity over time in downstream analyses, feature values at Day 1 were subtracted from Days 2 and 3 to create two datasets representing the net change in feature intensity over each time period (1 and 2 days, respectively). Notably, clustering analyses of these values do not show any clear patterns between donors or potency measures. (**Fig. 3A**). PCA of all samples from expansion days 1, 2 and 3 also showed no strong clustering corresponding to donor, but rather a pattern of samples clustering by the day of culture (**Fig. 3B**). Similar to the MS data analysis, linear regression was performed between each average spectral feature intensity and composite score for each MSC line. The top performing features (in terms of R^2^) are shown in **Fig. 3C, D**. Again, the changes in these individual spectral features showed reasonable correlation to the potency score, but after false discovery rate correction, these correlations were not significant. Particularly given the observed non-linear dynamics of many of these spectral features, it was hypothesized that identifying a panel of spectral features predictive of potency would be aided using diverse ML methods.

**Figure 3.**
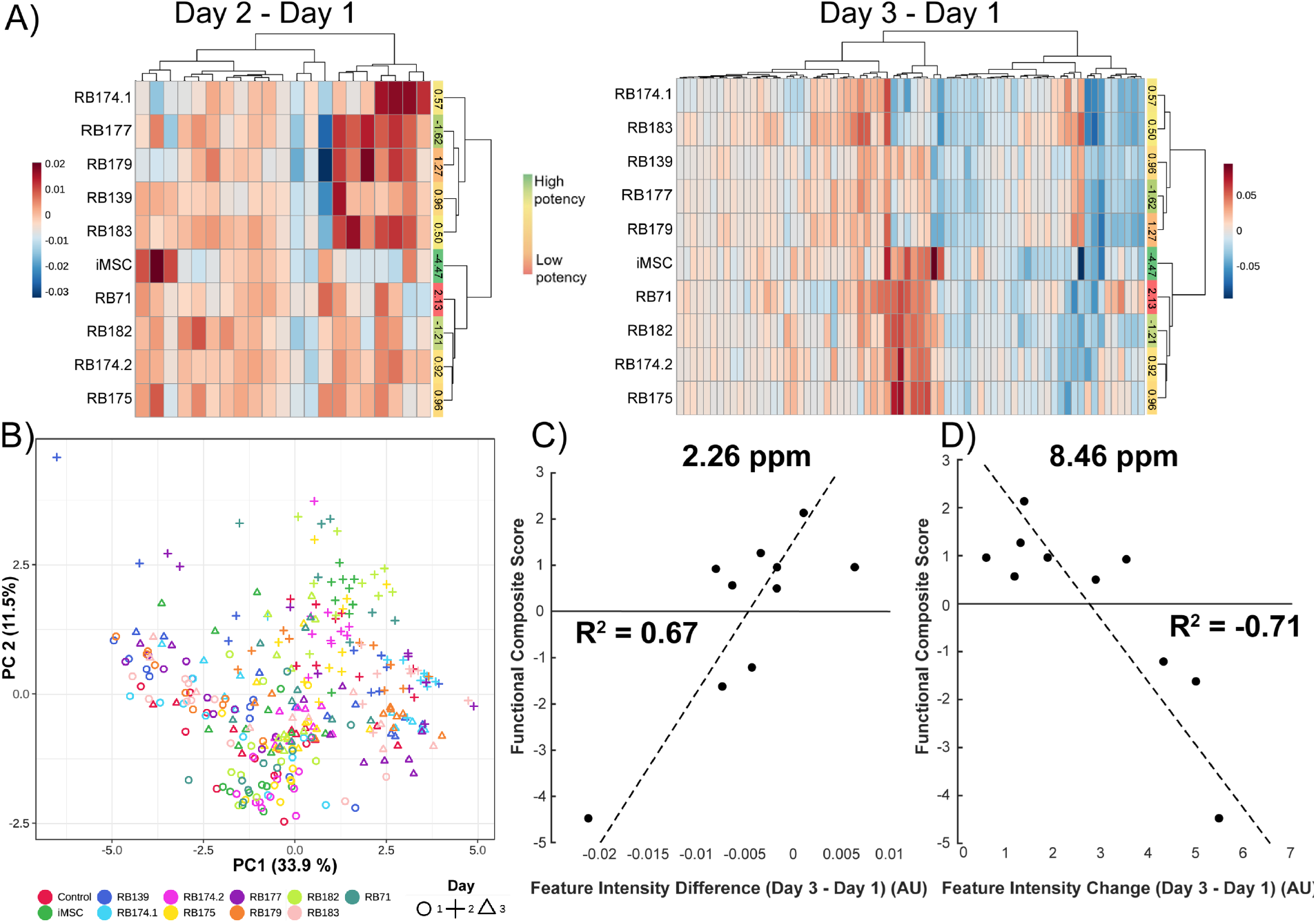
NMR analysis on daily media samples. A) Heatmap clustergram of Day 2-Day 1 highly variable feature intensities (21) and Day 3-Day 1 highly variable feature intensities (69). B) PCA scores plot of all samples from days 1-3, using all spectral features (138) as input. n=10 datapoints for each cell-line/timepoint with each cell-line represented by different color and each day by different shape. C) Regression of average donor Day 3 – Day 1 differences of feature at 2.26 ppm with eomposite functional score. D) Regression of average donor Day 3 – Day 1 differences of feature at 8.46 ppm with composite functional score.

### Modeling Approach and Identification of Consensus Predictive Metabolites

#### Evaluation of machine learning models

To find potency-related patterns in the data, several ML regression types were used for potency modeling (**Table 1**). Most models had comparable performance in both NMR and MS datasets based on LOO-R2, which indicates there are potency-related metabolic differences. This is reflected in metabolite abundances both intracellularly and in the cell media. Because the annotated metabolites and metabolite features identified in each model were not identical, we sought to develop a consensus modeling strategy to ensure metabolites of interest were robust and not unique to a particular ML approach. This strategy consisted of selecting only metabolites or features present in more than one of the initial models within a particular dataset – MS lipids, MS small polars (MS metabolite panel in **Supporting Information Table 4A, B**), NMR Day 2, or NMR Day 3 models were included in the consensus models. Most of the consensus models have comparable performance to the initial non-consensus models (in terms of LOO-R^2^), which underscores the robustness and predictive value of our consensus metabolite panels. To confirm these consensus models were not highly dependent on the iMSC sample, the final consensus metabolite panels were used to create another set of consensus models built and cross-validated on only the nine bone-marrow derived lines. The performance of these models, evaluated using LOO-R^2^, was comparable for MS models (LOO-R^2^ range from 0.72 to 0.99) while removal of iMSC from NMR consensus models resulted in lower LOO-R^2^ for most models (PLSR, SVM, GBR, LASSO, RF and DT). However, toe NMR consensus model constructed using symbolic regression was highly predictive of MSC potency (LOO-R^2^=0.96).

**Table 1.** Summary of Machine Learning Models to Predict MSC Potency. Machine learning (ML) models using various regression types – symbolic regression (SR), partial-least-squares regression (PLSR), support vector machine (SVM), gradient-boosted regression (GBR), least absolute shrinkage and selection operator (LASSO), random forest (RF), and decision tree (DT). Models created using four different input datasets: MS lipids, MS metabolites, NMR Day 3 – Day 1, and NMR Day 2 – Day 1 feature abundances. Consensus models created from only metabolites present in more than one of the initial models for both intracellular and extracellular metabolite datasets (using all 10 MSC lines). Final panels for the consensus models were used to create additional models trained and cross-validated on only the bone marrow derived lines (Consensus w/o iMSC, 9 total MSC lines).

| Input dataset | ML Regression Model - LOO-$R^2$ | | | | | | |
| --- | --- | --- | --- | --- | --- | --- | --- |
| Intracellular | SR | PLSR | SVM | GBR | LASSO | RF | DT |
| MS Lipids | 0.98 | 0.90 | 0.98 | 0.87 | 1.00 | 0.78 | 0.84 |
| MS Small Polars | 0.99 | 0.89 | 0.96 | 0.88 | 0.83 | 0.79 | 0.84 |
| Consensus | 0.99 | 0.89 | 1.00 | 0.88 | 0.92 | 0.83 | 0.60 |
| Consensus w/o iMSC | 0.99 | 0.94 | 0.89 | 0.86 | 0.88 | 0.80 | 0.72 |
| Extracellular | SR | PLSR | SVM | GBR | LASSO | RF | DT |
| NMR Day 3 – Day 1 | 0.99 | 0.90 | 0.99 | 0.88 | 0.93 | 0.71 | 0.91 |
| NMR Day 2 – Day 1 | 0.99 | 0.53 | 0.96 | 0.85 | 0.57 | 0.71 | 0.86 |
| Consensus | 0.99 | 0.87 | 0.82 | 0.89 | 0.72 | 0.60 | 0.86 |
| Consensus w/o iMSC | 0.96 | 0.64 | 0.42 | 0.75 | 0.31 | 0.50 | 0.68 |

#### Intracellular Metabolite Class Patterns in Modeling Results

Phosphatidylcholines (PC), phosphatidylethanolamines (PE) and sphingomyelins (SM) make up a large portion of the annotated dataset and show high importance in the models (**Fig. 4 A, B**). Interestingly, ether-linked phosphatidylethanolamines (PE-O) made up a smaller portion of the overall dataset compared to other lipid classes but were still among the top contributors to the models. Based on the weights applied to these lipid abundances in the regression models, they were highly important for model building. The over-representation of them in the models compared to the overall dataset is evidence that their abundances are related to MSC potency as measured by the composite score. This indicates a possible role in MSC functionality for members of this lipid class.

**Figure 4.**
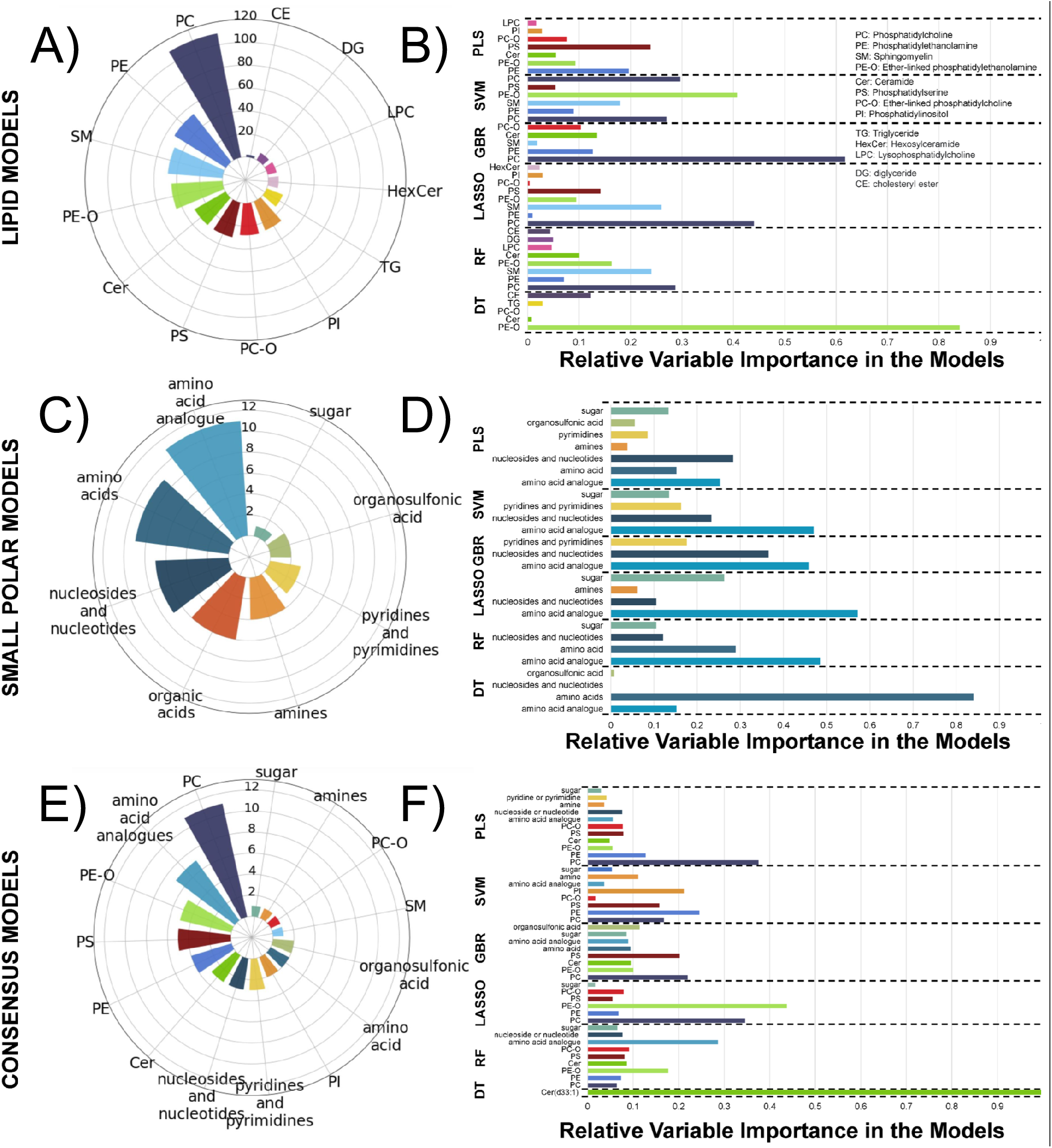
Mass Spectrometry Models to Predict MSC Potency. Radar plot (A) displaying the number of detected lipids in the annotated dataset organized according to lipid class. Bar plot (B) displaying the relative variable importance, calculated using the variable weights in the models, of each lipid class in each model type. Radar plot (C) displaying the number ob detected small polar metabolites in the annotated dataset organized according to class. Bar plot (D) dis playing the relative variable importance, calculated using the variable weights in the models, of molecule class in each model type. Radar plot (E) displaying the numbers of metabolites that presented in more than one initial model organized according to class. This list of consensus metabolites was used to create the consensus models. Bar plot (F) displaying the relative variable importance, calculated using the variable weights in the models, of each metabolite in the consensus models.

Amino acids and their analogues make up the majority of the small polar annotated dataset and showed importance in regression models (**Fig. 4 C, D**). Similar to PC in the lipid dataset, as they make up the largest portion of the data and contribute the most significantly to the models, no strong conclusions based on over-representation can be made. A hexose was detected in this dataset that was in the final panel for several of the models. However, there was only one sugar in the final annotated dataset meaning that class coverage for sugars was low. Given this finding, no strong conclusions about biological role or over-representation should be made.

The consensus metabolite list was made up of all metabolites that were in the final panel, following variable selection, of more than one initial model – lipid or small polar. This consensus list (**Supporting Information Table 4A,B**) was primarily composed of PC, amino acids and analogues, PE-O, and phosphatidylserines (PS) (**Fig. 4E**). These 41 metabolites from 15 classes (**Fig. 4E**) all showed importance in the models and were later investigated in pathway analysis. The variable presence plots for SR (**Supporting Information Figure S4**) indicate the following metabolites as having high occurrence in the final suite of models used for SR: acetyllysine, glucose isomer, hydroxykynurenamine, PE (O-32:1), PC(31:0), PE (O-38:2), and PI (36:2). In terms of the decision tree (DT) model, there was one ceramide (Cer(d33:1)) that was the sole contributor (**Fig. 4F**). Small changes in the data can cause large differences in DT models, making them relatively unstable compared to other types of models. It was also observed in this workflow that the DT regression models often selected a small number of predictive features. Since this particular lipid was not significantly important in other models as well, the large importance in the DT regression model may or may not be significant.

#### Different Consensus Media Metabolites Exhibit Distinct Changes

A total of 23 unique spectral features were selected as consensus features from both NMR timepoint datasets having been in top important features for at least two ML methods (**Fig. 5A, B**). Two features were common to both timepoint datasets (5.53 ppm and 5.30 ppm). The average trajectory over time for each of these consensus features is shown in **Fig. 5C**. As noted previously, many of these features show non-linear and non-monotonic behavior over time, and different changes between Day 1 and Day 2 or Day 3, accounting for some of the different features that were selected as consensus between the two datasets. Of the selected consensus features from both Day 3 and Day 2 NMR features, several were annotated to metabolites including proline, arginine, fructose, phenylalanine, pyruvate and an unknown uridine diphosphate-sugar (**Fig. 5D**). Of these metabolites, pyruvate and proline had some of the highest variable presence scores in the SR suite of models (**Supporting Information Figure S4**) as well as several unknown, unannotated metabolites. Several features that appeared as important in some ML methods were not matched to spectral databases and may require further experimentation to confidently annotate. Particularly, two of these unknown features at 5.53 ppm and 5.30 ppm appeared as consensus features in both timepoint datasets, suggesting that whatever metabolite(s) these features correspond to may be a robust predictor of potency across time.

**Figure 5.**
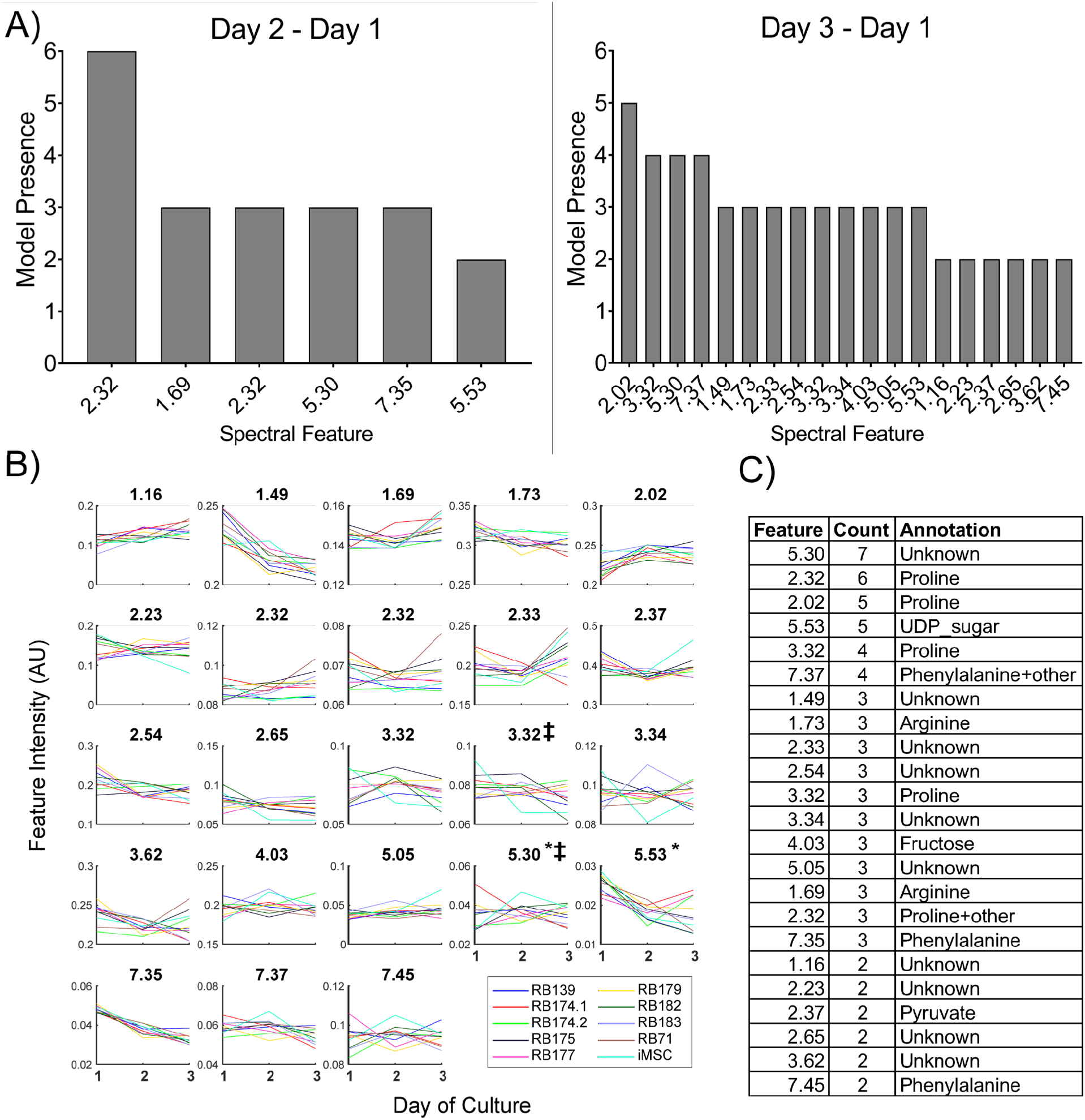
Consensus NMR metabolite feature trends and annotation. A) Consensus features selected as important across all modeling methods for Day 3-Day 1, and Day 2 – Day 1 datasets. Counts indicate for how many models each feature was selected within top 10% of important variables for prediction. Names of features represent approximate ppm of quantified spectral peak. B) Average spectral feature intensity trajectories over Days 1-3 (n=10 per donor per timepoint). Feature names indicate approximate chemical shift values of integrated peak. Intensity values are in arbitrary units. * Indicates consensus features in both timepoint detasets. ‡ indicates features identified from SR consensus model. C) Putative metabolite annotations of consensus spectral features. UDP = uridine diphosphate

### Interpretation of Consensus Metabolites Through Pathway Analysis

The consensus list from each modeling set (**Table S4A, B** for MS and **Fig. 5D** for NMR) was used to search for potency-associated changes in metabolism on the pathway level. Of the top enriched pathways, there were 7 with significant p-values (p<0.05) for the NMR consensus metabolites (**Fig. 6A**), 12 with significant p-values (p<0.05) for MS lipidomic analysis (**Fig. 6B**), and none for MS small polar analysis (**Fig. 6C**). Top lipid enriched pathways included sphingolipid signaling pathway, autophagy, and necroptosis, indicating that these lipids could be important in MSCs based on their role in cell cycle. Approximately 10% of the small polar dataset was able to be annotated, compared to the lipid dataset which was 45% annotated. Because enrichment analysis relies on over-representation of metabolites in particular pathways, the lower metabolite coverage of the small polar dataset was likely a contributing factor to the lack of significance of the pathways identified. Additionally, some of the identified metabolites in this dataset were not found in pathways in the Small Molecule Pathway Database used for pathway enrichment. Despite the lack of significantly enriched pathways in the MS metabolite data, interestingly there was some agreement between top resulting pathways in the NMR and MS small polar analyses. In particular, aspartate metabolism, arginine-proline metabolism, glycine-serine metabolism, and glucose-alanine metabolism appeared in the top enriched pathways for both datasets. This points to some potential consistency in similar metabolic pathways responsible for predicting MSC potency both early in-process and at end-process. This points to some potential consistency in similar metabolic pathways responsible for predicting MSC potency both early in-process and at end-process.

**Figure 6.**
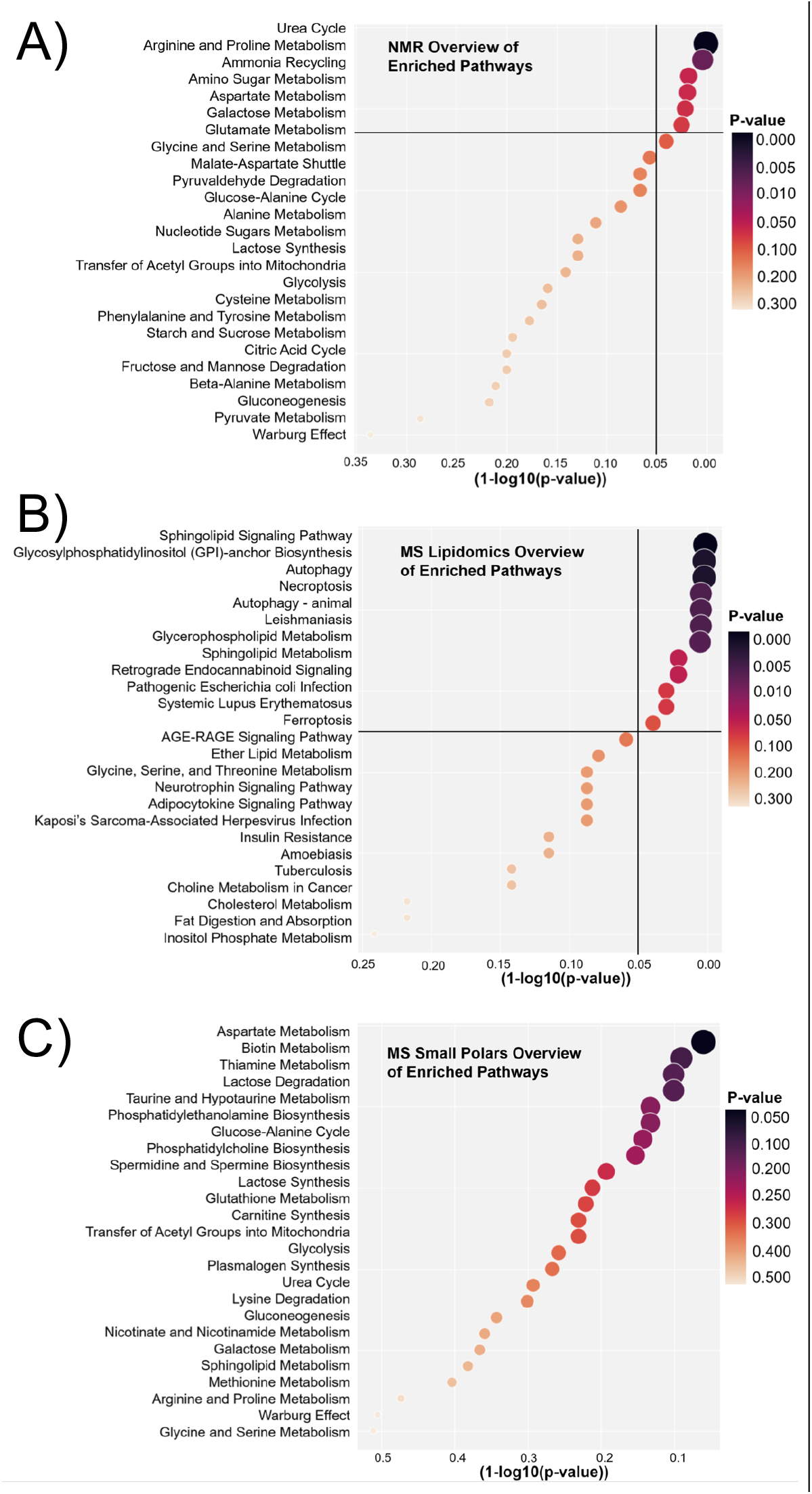
Enriched pathways identified from MS and NMR consensus metabolite datasets. Pathway enrichments plots from NMR metabolite modeling consensus list with pathways and p-values calculated using Metaboanalyst (A), MS lipid modeling consensus list with pathways and p-values calculated using LIPEA (B), and MS small polar consensus list with pathways and p-values calculated using Metaboanalyst (C). Size and color of markers scaled according to p-value.

## DISCUSSION

The identification of CQAs correlative to immunomodulatory potency would enable predictive approaches to address some of the grand challenges that hinder the approval and clinical use of MSCs as cell therapies.^16,17^ Donor-donor variability and different ex vivo manufacturing procedures create inconsistencies in the therapeutic potency of MSCs, and ultimately efficacy outcomes when evaluated in clinical trials.^12,13^ In this study, we greatly expanded on our previous work^26^ through incorporation of additional cell-lines, assessment of in-process and end-process metabolites, and development of comprehensive ML modeling approach to identify predictive markers. The in-process media analysis might inform decisions made early in cell expansion phase that if translated to a manufacturing setting would reduce manufacturing cost due to identification of failed batches early in manufacturing. For the broad-spectrum analysis of intracellular metabolomics, the discovery of small molecules and lipids correlative to MSC immunomodulation, as a panel of CQAs, is of great interest and a means to distinguish biological heterogeneity and predict the in vivo therapeutic potency of the MSC product. Moreover, a composite score indicative of immunomodulatory potency was developed based on cumulative results from multiple in vitro potency assays to enable correlations of top features from media and intracellular metabolome. This robust approach accounts for the variability in MSC functional responses and identifying consensus top features, i.e. potential CQAs, using ML models.

An inherent limitation of relying on a single ML model to inform decisions or hypotheses about data can be intrinsic bias based on the specific framework and assumptions that go into using a specific method.^39,40^ By using a diverse array of ML regression methods, we avoid being biased too strongly by a single method in identifying features and metabolites predictive of MSC potency. Our consensus approach to identifying potential CQAs reduces the possibility of model specific results by ensuring that they are deemed important by multiple ML methods as illustrated previously.^25^ Ultimately the biggest limitation to this method is the size of our dataset, which is limited by the amount of cell material needed for both functional assays and analytical measurements. However, we attempt to mitigate the impact of overfitting on the interpretation of our results using cross-validated model tuning and leave-one-out model performance calculations, in addition to performing our ML analysis without our “outlier” iMSC donor to assess that unique group’s impact on our results.

Lipid classes such as phosphatidylcholines (PC), phosphatidylethanolamines (PE), phosphatidylinositols (PI), and phosphatidylserines (PS) are all glycerophospholipids that were found as predictors in our ML models.^4141^ Differences in MSC glycerophospholipid composition has been shown between young and old MSC donors as well as early and late passage MSCs.^42,43^ PC and PE were two of the most abundant lipid classes found in the ML models and are two of the most abundant glycerophospholipids found in mammalian cells.^41^ PC account for roughly 50% of all cellular phospholipids, and have been shown to be predictive of MSC immunomodulation.^26,43,44^ The majority of PE are found in the mitochondrial membrane, and MSC mitochondrial fitness is associated with its glycolytic potential.^23,44,45^ Greater glycolytic potential has been shown to be associated with greater MSC immunomodulation.^23,45^ PE have also been shown to positively regulate autophagy, which helps prevent cellular ageing.^46^ Several studies have shown that increased autophagy in MSCs can help prevent senescence, increase survival and engraftment, and increase immunomodulatory function.^47,48^ As mentioned previously, PI were also predictors of MSC immunomodulation and are precursors for the biosynthesis of glycosylphosphatidylinositol (GPI) anchors.^49^ GPI anchored markers such as CD157, which aids in immunomodulation, is involved in migration, self-renewal, osteogenic differentiation, and mitochondrial transfer in MSCs.^50–52^ Another glycerophospholipid, PS, is an important molecule in apoptosis signaling, and *in vivo* studies have suggested that MSC apoptosis may be crucial for their therapeutic efficacy.^53–55^ Two other lipid classes found as predictors, sphingomyelins (SM) and ceramides (Cer), are closely related to one another via the sphingolipid metabolic pathway.^56^ Increases in sphingomyelin from ceramide treatment has been shown to increase senescence in bone marrow MSC (BMMSC)s.^57^ Increased levels of acyl chain ceramides are also associated with decreased levels of IDO activity in BMMSCs.^56^ The sphingolipid signaling pathway, a significant pathway found in our results, also plays an important role in MSC migration and osteogenic differentiation.^58,59^

Similarly, we sought to identify metabolites in the media during expansion that relate to immunomodulatory function as this represents a non-destructive, in-process approach for monitoring cell quality. In-process monitoring allows for greater control and quality assurance of cell therapies throughout the expansion process.^60,61^ In-process monitoring of parameters such oxygen diffusion, CO2, pH, temperature, and glucose and lactate consumption/production are well established in biomanufacturing.^62^ New methods such as gas chromatography mass spectrometry have been used to measure biomarkers predictive of MSC immunomodulation.^63^ Here, we profiled media metabolites from the first three days of expansion to predict MSC function at the end of manufacturing. The amino acids proline, arginine, phenylalanine, and aspartate were all predictive of MSC immunomodulation. Arginine and proline metabolism has been associated with autophagy of MSCs, which was found as a significant pathway from our lipid analysis.^64^ Increased ammonia is a by-product of protein and amino acid catabolism and is converted to urea through the urea cycle with arginine and aspartate being key amino acids in the urea cycle.^65,66^ Aspartate metabolism is also associated with the TCA cycle which is increased from cellular OXPHOS with pyruvate being an intermediate of both OXPHOS and glycolysis.^31^ As mentioned previously, metabolic shifts in MSCs from glycolysis to OXPHOS is associated with a decrease in MSC immunomodulation.^23^ Similarly, IFN-γ and hypoxic conditioning increases glycolysis in MSCs and increases the capacity for glucose and fructose uptake.^67^ Additionally, arginine and proline metabolism, amino sugar metabolism, and galactose metabolism showed significant differences when comparing adipose-derived MSCs (ADMSCs) and BMMSCs.^68^ Upregulation of genes associated with galactose metabolism has also been associated with higher immunomodulation of MSCs.^69^ These results further emphasize the critical role of metabolism during MSC manufacturing and how our robust machine learning approach can identify pathways relevant to MSC therapeutic potential.

## CONCLUSION

Overall, this study establishes a comprehensive framework for future studies to interrogate metabolites as predictive markers for MSCs when changing manufacturing parameters. A major example of a significant manufacturing change would be increasing manufacturing scale for clinical trials as the average dose for MSC based therapies is on the order of 10^8^ cells for a single patient.^14^ Because of this, scaling up to bioreactors is necessary to produce enough cells for the clinic. Scaling up MSC manufacturing has a significant effect on parameters such as nutrient transport and MSC metabolism.^31^ Another parameter is the type of media used for the expansion of MSCs. Priming MSCs with inflammatory factors such as IFN-γ or TNF-α, as well as hypoxia, are also being explored for MSC therapies due to the potential of increasing their therapeutic potential.^23,67,70^ Priming conditions have significant effects on MSC metabolism and could potentially lead to greater therapeutic outcomes through a more homogeneous, potent MSC product.^23,31,67^ Lastly, MSC metabolism could further predict MSC engraftment and survivability *in vivo*, which together have been linked to clinical outcomes.^71^ By using multiple MSC lines and ML models, this study sets the framework for rigorous predictive marker identification that can be used in future studies to help address potential manufacturing process hurdles for MSC therapeutics. Based on the metabolite classes identified in this work (both in-process and at the end of expansion), targeted assays can be developed for better MSC potency assessment and release criteria for immunomodulatory therapeutics.

## Acknowledgements

We would like to thank Julie Nelson in the University of Georgia Flow Cytometry Core (Center for Tropical and Emerging Global Diseases) for providing her expertise with flow cytometry equipment. We thank the Systems Mass Spectrometry Core at the Georgia Institute of Technology for valuable support in LC-MS experiments.

