## Supporting Information for "Development of a Robust Consensus Modeling Approach for Identifying Cellular and Media Metabolites Predictive of Mesenchymal Stromal Cell Potency"

**Table S1. MSC donor and expansion information.** *indicates MSC lines used in previous study (ref26).

| **Donor** | **Sex** | **Age**  **(years)** | **Total cells seeded (C_i_)** | **Final cell yield (C_f_)** | **Initial population doubling level (PDL_0_)** | **Final PDL** |
| --- | --- | --- | --- | --- | --- | --- |
| RB139 | Male | 25 | 3.75E+05 | 8.10E+06 | 12.2 | 16.6 |
| RB174.1 | Male | 25 | 3.75E+05 | 9.80E+06 | 13.06 | 17.8 |
| RB177 | Male | 22 | 3.75E+05 | 4.23E+06 | 15.1 | 18.6 |
| RB179 | Male | 21 | 3.75E+05 | 8.20E+06 | 12.6 | 17.1 |
| RB183 | Female | 26 | 3.75E+05 | 5.30E+06 | 12 | 15.8 |
| RB71* | Female | 18-30 | 3.75E+05 | 7.50E+06 | 12.72 | 17.0 |
| RB174.2 | Male | 25 | 3.75E+05 | 8.43E+06 | 13.06 | 17.6 |
| RB175 | Male | 25 | 3.75E+05 | 6.24E+06 | 12 | 16.1 |
| RB182* | Female | 26 | 3.75E+05 | 3.65E+06 | 11.48 | 14.8 |
| iMSC* | Not available | | 3.75E+05 | 1.86E+07 | Not available | |

**Table S2.** Antibodies used for T cell suppression assay

| **Laser** | **EM Filter** | **Marker** | **Color / Format** | **Host / Target** | **Isotype** | **Clone** | **Company** | **Catalog** |
| --- | --- | --- | --- | --- | --- | --- | --- | --- |
| 405 | 710/50 | CD4 | Brilliant Violet 711 | Mouse anti-Human | IgG1 κ | RPA-T4 | BioLegend | 300558 |
| 638 | 660/20 | CD8 | APC | Mouse anti-Human | IgG1 | CN9V1 | BioLegend | orb248718 |
| 488 | 525/40 | CFSE | CFSE | N/A | N/A | N/A | BioLegend | 423801 |
| 405 | 610/20 | Zombie Yellow | Zombie Yellow | N/A anti-All Species | N/A | N/A | BioLegend | 423104 |

**Table S3. UHPLC chromatographic gradients.** Mobile Phase A used for HILIC chromatography was 80:20 water:MeCN with 10 mM ammonium formate and 0.1% formic acid. Mobile phase B for HILIC chromatography was acetonitrile with 0.1% formic acid. The flow rate was set at 0.4 mL/min.  The column temperature was set to 40 °C, the injection volume was 2 µL. Mobile Phase A for Reverse Phase chromatography in positive mode was 40:60 water: acetonitrile with 10 mM ammonium formate and 0.1% formic acid and Mobile Phase B was 10:90 acetonitrile:isopropyl alcohol, with 10 mM ammonium formate and 0.1% formic acid. For negative ionization mode, the mobile phases were 40:60 water:acetonitrile with 10 mM ammonium acetate (mobile phase A) and 10:90 acetonitrile:isopropyl alcohol, with 10 mM ammonium acetate (mobile phase B). The flow rate was set at 0.40 mL min^-1^. The column temperature was set to 50 °C, and the injection volume was 2 µL.

| **HILIC Chromatography** | | **Reverse Phase Chromatography** | |
| --- | --- | --- | --- |
| 0-8 min | 5% Mobile Phase A | 0-1 min | 80% Mobile Phase A |
| 8-10.4 min | 60% Mobile Phase A | 1-5 min | 40% Mobile Phase A |
| 10.5-14 min | 5% Mobile Phase A | 5-5.5 min | 30% Mobile Phase A |
|  |  | 5.5-8 min | 15% Mobile Phase A |
|  |  | 8-8.2 min | 10% Mobile Phase A |
|  |  | 8.2-10.5 min | 0% Mobile Phase A |
|  |  | 10.7-12 min | 80% Mobile Phase A |

**Table S4.** **Proposed annotations for metabolites in models**. The table shows high resolution experimental *m/z* value for the species of interest, proposed annotation, main adduct type detected, retention time and metabolite ID which includes which chromatography type and ionization mode, elemental formula, mass error (ppm) calculated from exact monoisotopic mass, annotation confidence level, MS/MS collision energies, and the online database used for the MS^2^ spectra library match. The confidence level for lipid annotation was assigned as (1) exact mass, isotopic pattern, and MS/MS spectrum of a chemical standard matched to the feature. (2) exact mass, isotopic pattern, retention time, and MS/MS spectrum matched to an in-house spectral database or literature spectra (3) putative ID assignment based only on elemental formula match. (4) unknown compound. Asterisks (*) designate compounds for which MS^2^ spectral match was done using in-silico fragmentation in Compound Discoverer.

A)
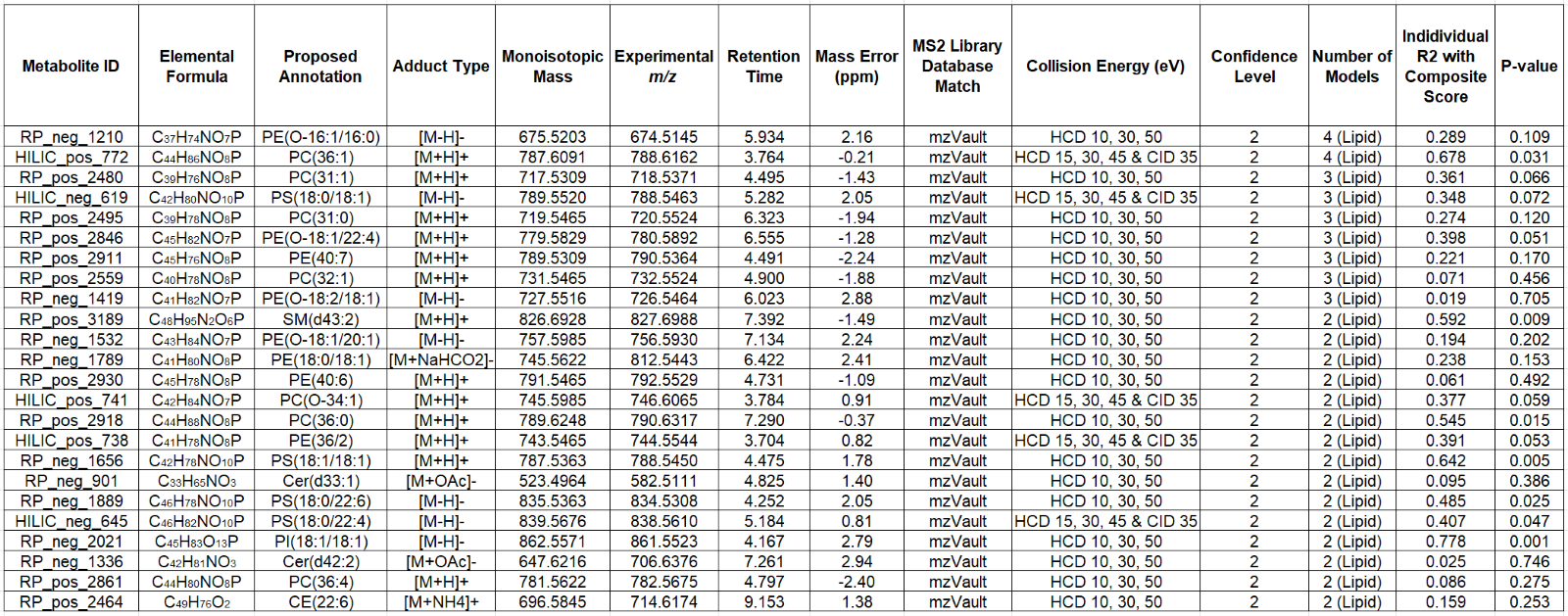
Proposed lipid annotations for all lipids that presented in the final panel for more than one lipid model
B)
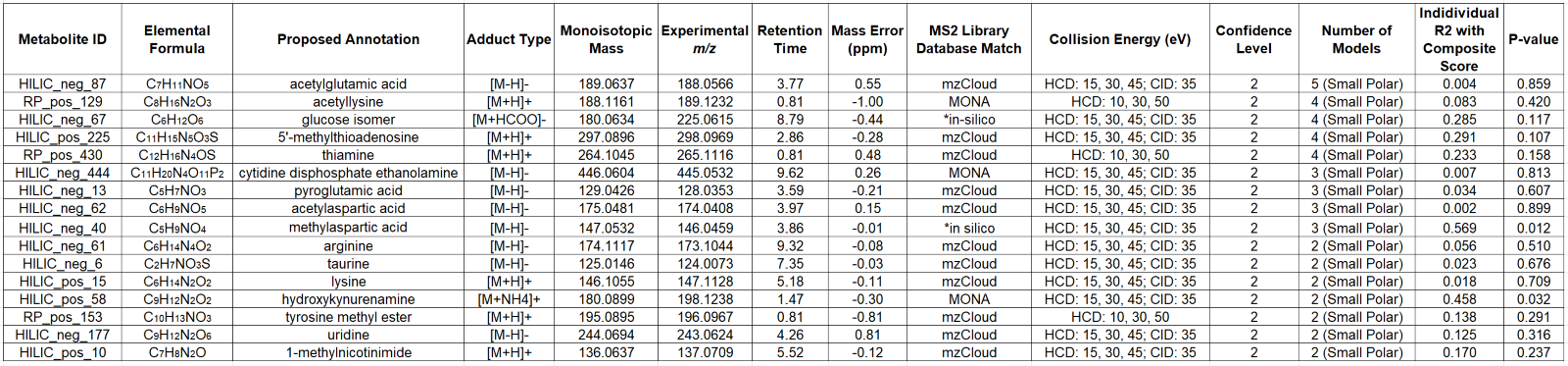
Proposed metabolite annotations for all metabolites that presented in the final panel for more than one small polar model. *in-silico fragmentation in Compound Discoverer.
C)
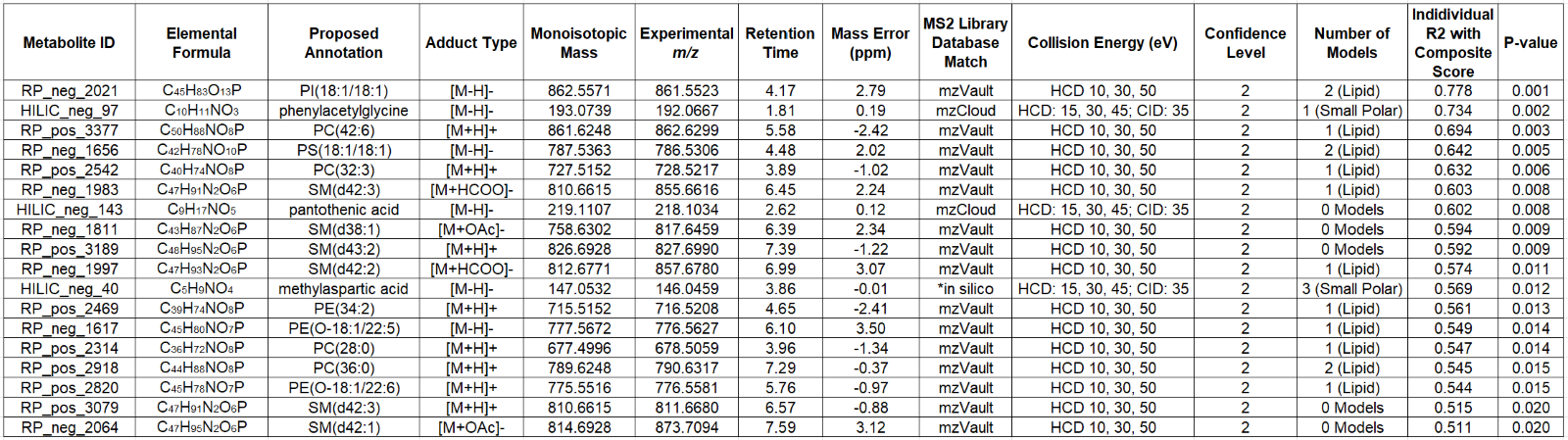
All metabolites, regardless of class, that had individual correlations with potency score with an R^2^ of 0.5 or greater.

**SUPPLEMENTAL FIGURES**

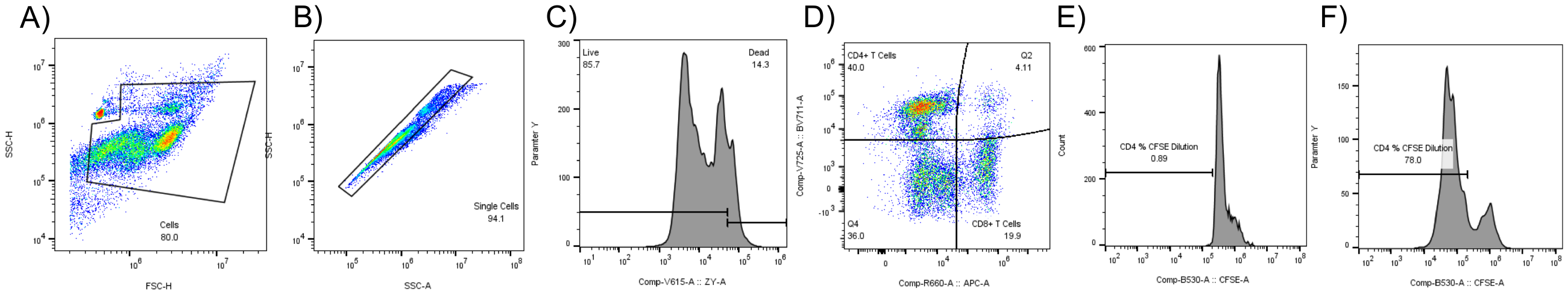

**Figure S1. Gating strategy to assess T cell proliferation.** Cellular debris and Dynabeads were first removed (A) followed by cell doublets (B). Using FMO controls, live cells were then gated based on Zombie Yellow staining (C) and used to determine CD4+ and CD8+ T cell populations (D). Negative control PBMCs (No stimulation, no MSCs) were then used for the CFSE dilution gate (E) and positive control PBMCs (stimulation, no MSCs) were used for baseline comparison (F).

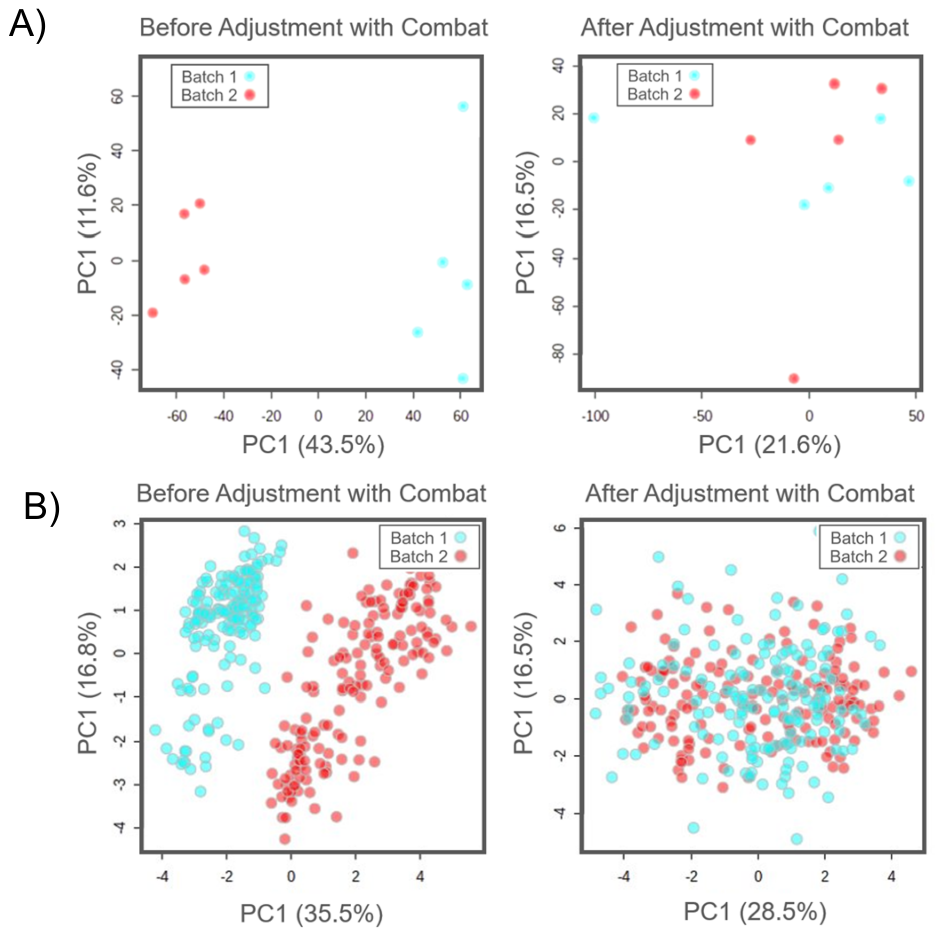

**Figure S2. Batch Correction of MS and NMR datasets.** MS (A) and NMR (B) results of PCA scores plot before and after batch adjustment using Combat in Metaboanalyst. For MS, each point is one of the cell lines at end-of-process. For NMR, each point is a sample from each culture replicate at each timepoint.

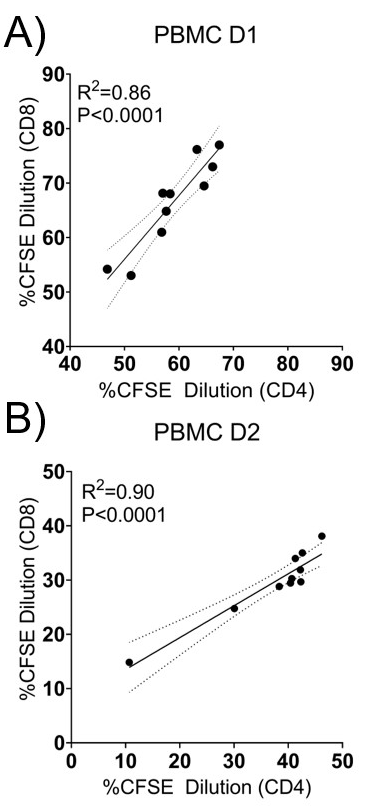

**Figure S3. Correlation of CD4 and CD8 T cell proliferation.** Linear Regression of CD4/CD8 Proliferation for PBMC donor 1 (A) and PBMC donor 2 (B)

**Figure S4: Evolved Analytics Data Modeler ML consensus feature results.** Pareto front plot of final suite of models using consensus features for MS (A) and NMR ( C). Bar plots of metabolites from consensus feature list present in the final suite of models for MS (B), and NMR (D).
